## Supplementary Figures 1-8 for "Large-scale animal model study uncovers altered brain pH and lactate levels as a transdiagnostic endophenotype of neuropsychiatric disorders involving cognitive impairment"

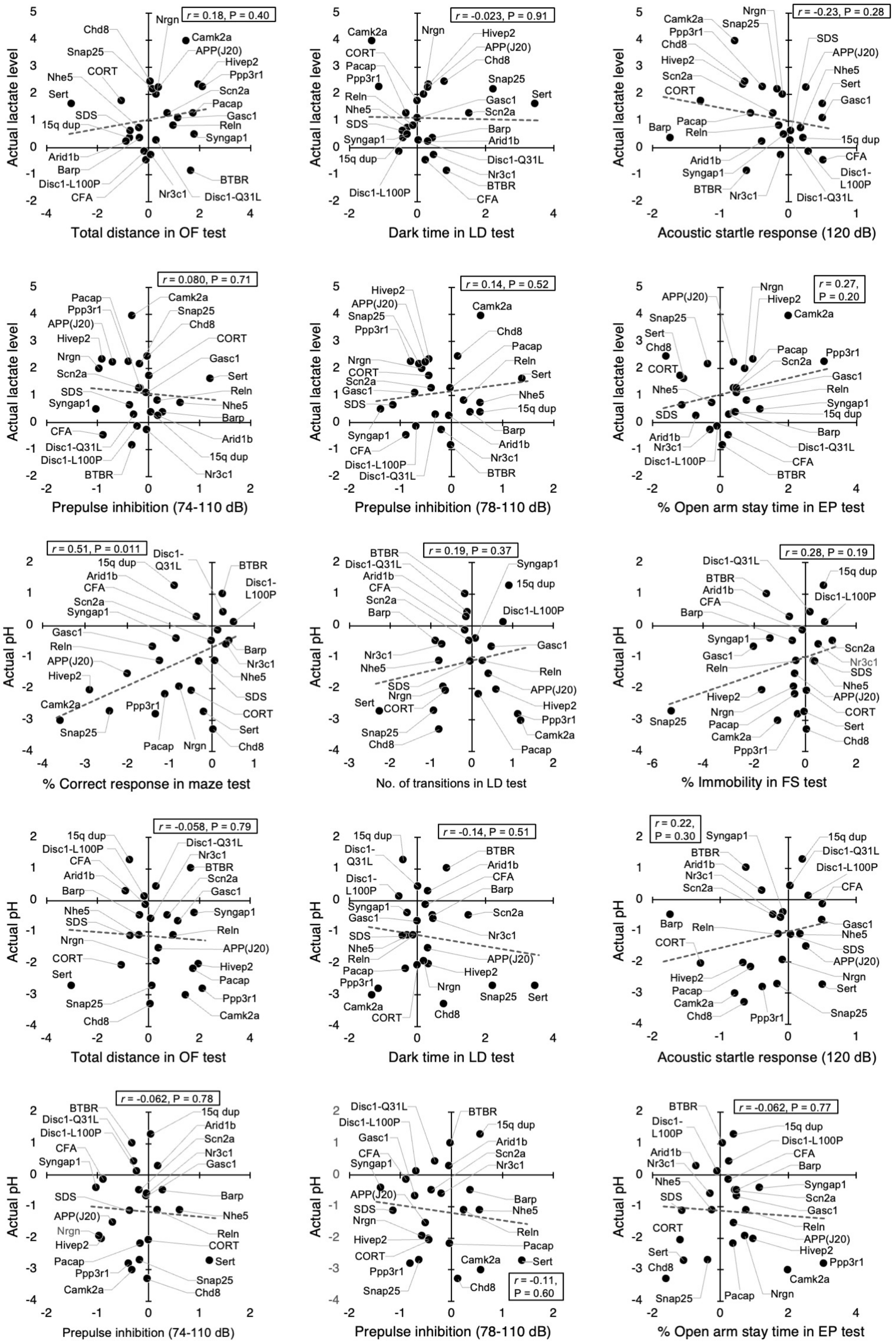
r = 0.18, P = 0.40
Actual lactate level
Total distance in OF test
r = -0.023, P = 0.91
Actual lactate level
Dark time in LD test
r = -0.23, P = 0.28
Actual lactate level
Acoustic startle response (120 dB)
r = 0.080, P = 0.71
Actual lactate level
Prepulse inhibition (74-110 dB)
r = 0.14, P = 0.52
Actual lactate level
Prepulse inhibition (78-110 dB)
r = 0.27, P = 0.20
Actual lactate level
% Open arm stay time in EP test
r = 0.51, P = 0.011
Actual pH
% Correct response in maze test
r = 0.19, P = 0.37
Actual pH
No. of transitions in LD test
r = 0.28, P = 0.19
Actual pH
% Immobility in FS test
r = -0.058, P = 0.79
Actual pH
Total distance in OF test
r = -0.14, P = 0.51
Actual pH
Dark time in LD test
r = 0.22, P = 0.30
Actual pH
Acoustic startle response (120 dB)
r = -0.062, P = 0.78
Actual pH
Prepulse inhibition (74-110 dB)
r = -0.11, P = 0.60
Actual pH
Prepulse inhibition (78-110 dB)
r = -0.062, P = 0.77
Actual pH
% Open arm stay time in EP test

**Figure S1. Correlations of brain lactate levels and pH with behavioral measures in an exploratory cohort.** Scatter plots showing effect size-based correlations between actual lactate levels and pH, and behavioral measures. Data from 24 strains/conditions of mice used in the prediction analysis are shown. EP, elevated-plus maze; FS, forced swim test; LD, light/dark transition test; OF, open field test; *r*, Pearson's correlation coefficient.

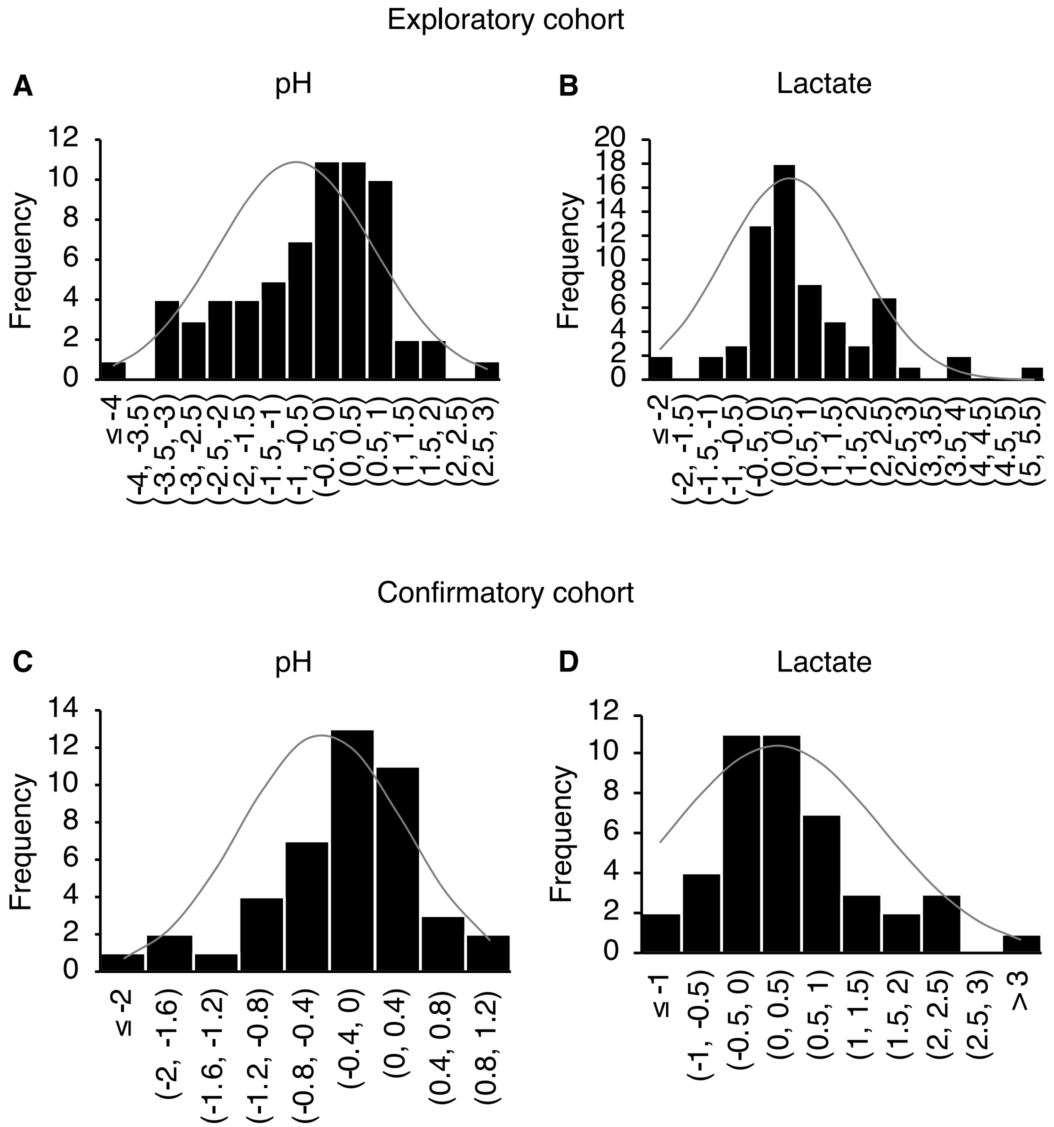


**Figure S2. Normal distribution of effect size values for pH and lactate in the exploratory and confirmatory cohorts.** (A) D = 0.12, p = 0.32. (B) D = 0.15, P = 0.088. (C) D = 0.14, P = 0.33. (D) D = 0.18, P = 0.10.

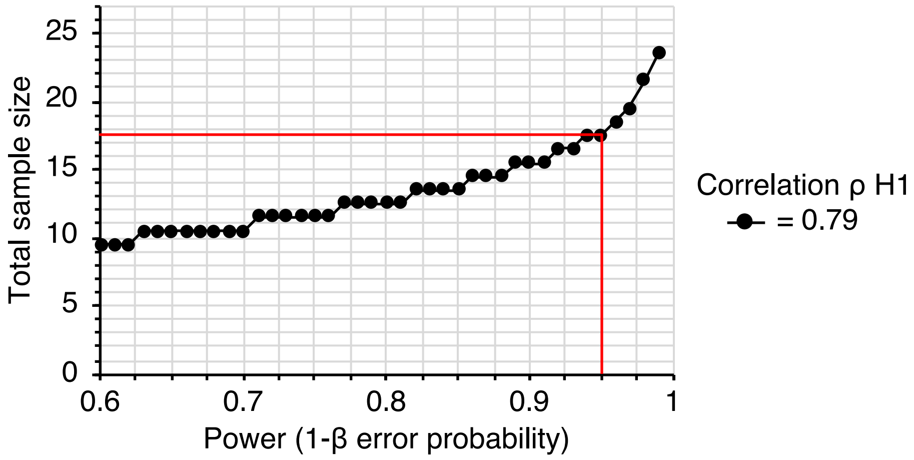


**Figure S3. A *priori* power analysis to estimate the optimum sample size for the confirmatory experiment.** Input parameters: tails = two, correlation |ρ| H1 = 0.79, α error probability = 0.01, power (1–β error probability) = 0.95, correlation |ρ| H0 = 0. Output parameters: total sample size = 18, actual power = 0.95. The red line indicates 1–β = 0.95.

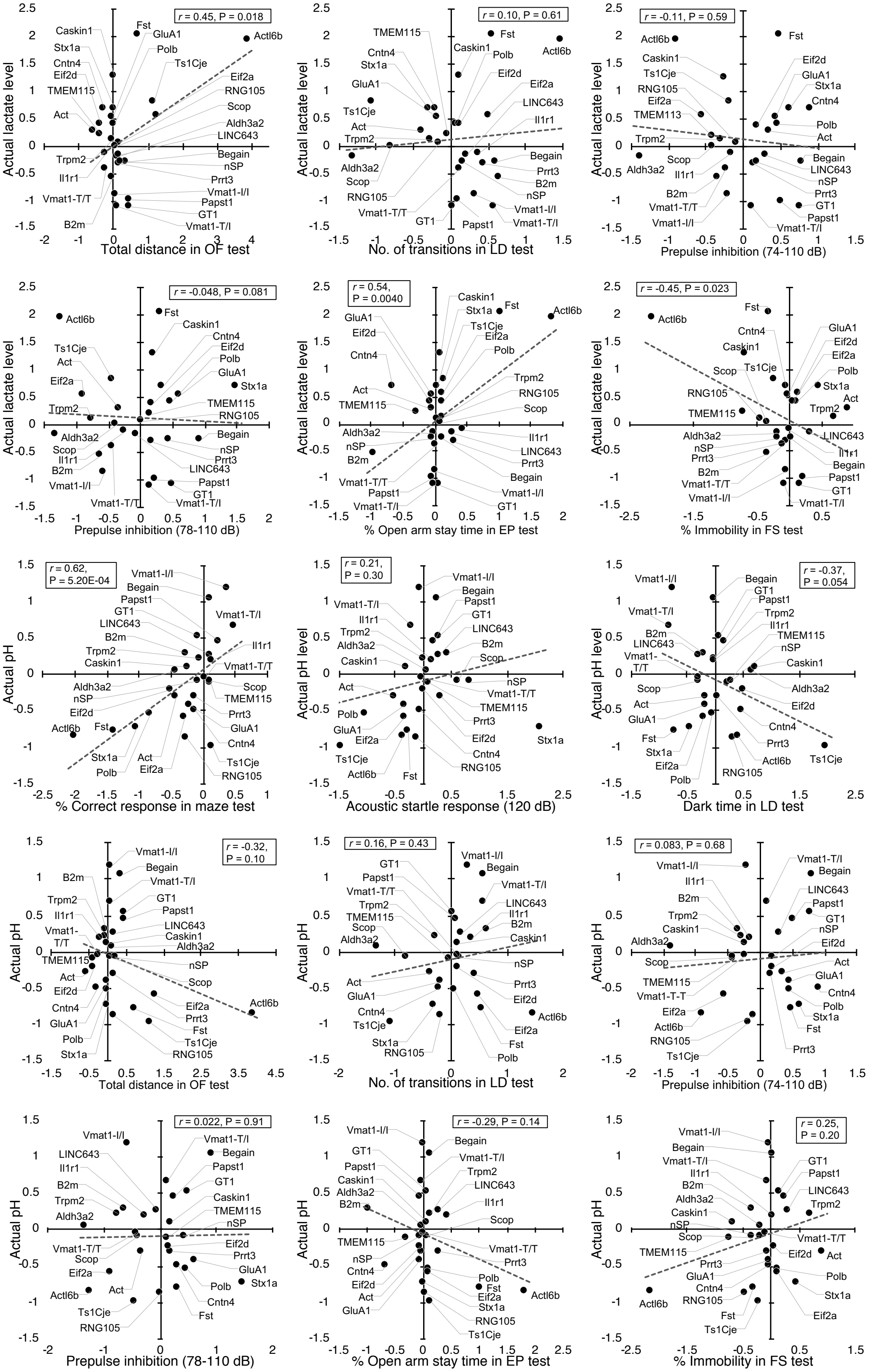
Actual lactate level
Total distance in OF test
r = 0.45, P = 0.018
Caskin1
Stx1a
Cntn4
Eif2d
TMEM115
Act
Trpm2
Il1r1
Vmat1-T/T
B2m
Fst
GluA1
Polb
Ts1Cje
Actl6b
Eif2a
RNG105
Scop
Aldh3a2
LINC643
Begain
nSP
Prrt3
Vmat1-I/I
Papst1
GT1
Vmat1-T/I
No. of transitions in LD test
r = 0.10, P = 0.61
TMEM115
Cntn4
Stx1a
GluA1
Ts1Cje
Act
Trpm2
Aldh3a2
Scop
RNG105
Vmat1-T/T
GT1
Caskin1
Fst
Polb
Actl6b
Eif2d
Eif2a
LINC643
Il1r1
Begain
Prrt3
B2m
nSP
Vmat1-I/I
Papst1
Vmat1-T/I
Prepulse inhibition (74-110 dB)
r = -0.11, P = 0.59
Actl6b
Caskin1
Ts1Cje
RNG105
Eif2a
TMEM113
Trpm2
Aldh3a2
Scop
Il1r1
B2m
Vmat1-T/T
Vmat1-I/I
Fst
Eif2d
GluA1
Stx1a
Cntn4
Polb
Act
Begain
LINC643
nSP
Prrt3
GT1
Papst1
Vmat1-T/I
Prepulse inhibition (78-110 dB)
r = -0.048, P = 0.081
Actl6b
Ts1Cje
Act
Eif2a
Trpm2
Aldh3a2
Scop
Il1r1
B2m
Vmat1-I/I
Vmat1-T/T
Fst
Caskin1
Cntn4
Eif2d
Polb
GluA1
Stx1a
TMEM115
RNG105
Begain
nSP
Prrt3
LINC643
Papst1
GT1
Vmat1-T/I
% Open arm stay time in EP test
r = 0.54, P = 0.0040
GluA1
Eif2d
Cntn4
Act
TMEM115
Aldh3a2
nSP
B2m
Vmat1-T/T
Papst1
Vmat1-T/I
Caskin1
Stx1a
Fst
Ts1Cje
Eif2a
Polb
Actl6b
Trpm2
RNG105
Scop
Il1r1
LINC643
Prrt3
Begain
Vmat1-I/I
GT1
% Immobility in FS test
r = -0.45, P = 0.023
Actl6b
Fst
Cntn4
Caskin1
Scop
Ts1Cje
RNG105
TMEM115
Aldh3a2
nSP
Prrt3
B2m
Vmat1-T/T
Vmat1-I/I
GluA1
Eif2d
Eif2a
Polb
Stx1a
Act
Trpm2
LINC643
Il1r1
Begain
Papst1
GT1
Vmat1-T/I
Actual pH
% Correct response in maze test
r = 0.62, P = 5.20E-04
Vmat1-I/I
Begain
Papst1
GT1
LINC643
B2m
Trpm2
Caskin1
Aldh3a2
nSP
Eif2d
Actl6b
Fst
Stx1a
Polb
Act
Eif2a
Vmat1-T/I
Il1r1
Vmat1-T/T
Scop
TMEM115
Prrt3
GluA1
Cntn4
Ts1Cje
RNG105
Actual pH level
Acoustic startle response (120 dB)
r = 0.21, P = 0.30
Vmat1-T/I
Il1r1
Trpm2
Aldh3a2
Caskin1
Act
Polb
GluA1
Eif2a
Ts1Cje
Actl6b
Fst
Vmat1-I/I
Begain
Papst1
GT1
LINC643
B2m
Scop
nSP
Vmat1-T/T
TMEM115
Prrt3
Eif2d
Cntn4
RNG105
Stx1a
Dark time in LD test
r = -0.37, P = 0.054
Vmat1-I/I
Vmat1-T/I
B2m
LINC643
Vmat1-T/T
Scop
Act
GluA1
Fst
Stx1a
Eif2a
Polb
Begain
GT1
Papst1
Trpm2
Il1r1
TMEM115
nSP
Caskin1
Aldh3a2
Eif2d
Cntn4
Prrt3
Actl6b
RNG105
Ts1Cje
Total distance in OF test
r = -0.32, P = 0.10
B2m
Trpm2
Il1r1
Vmat1-T/T
TMEM115
Act
Eif2d
Cntn4
GluA1
Polb
Stx1a
Vmat1-I/I
Begain
Vmat1-T/I
GT1
Papst1
LINC643
Caskin1
Aldh3a2
nSP
Scop
Actl6b
Eif2a
Prrt3
Fst
Ts1Cje
RNG105
No. of transitions in LD test
r = 0.16, P = 0.43
GT1
Papst1
Vmat1-T/T
Trpm2
TMEM115
Scop
Aldh3a2
Act
GluA1
Cntn4
Ts1Cje
Stx1a
RNG105
Vmat1-I/I
Begain
Vmat1-T/I
LINC643
Il1r1
B2m
Caskin1
nSP
Prrt3
Eif2d
Actl6b
Eif2a
Fst
Polb
Prepulse inhibition (74-110 dB)
r = 0.083, P = 0.68
Vmat1-I/I
Il1r1
B2m
Trpm2
Caskin1
Aldh3a2
Scop
TMEM115
Vmat1-T-T
Eif2a
Actl6b
RNG105
Ts1Cje
Vmat1-T/I
Begain
LINC643
Papst1
GT1
nSP
Eif2d
Act
GluA1
Cntn4
Polb
Stx1a
Fst
Prrt3
Prepulse inhibition (78-110 dB)
r = 0.022, P = 0.91
Vmat1-I/I
LINC643
Il1r1
B2m
Trpm2
Aldh3a2
Vmat1-T/T
Scop
Eif2a
Actl6b
Act
Ts1Cje
RNG105
Vmat1-T/I
Begain
Papst1
GT1
Caskin1
TMEM115
nSP
Eif2d
Prrt3
GluA1
Stx1a
Polb
Cntn4
Fst
% Open arm stay time in EP test
r = -0.29, P = 0.14
Vmat1-I/I
GT1
Papst1
Caskin1
Aldh3a2
B2m
TMEM115
nSP
Cntn4
Eif2d
Act
GluA1
Begain
Vmat1-T/I
Trpm2
LINC643
Il1r1
Scop
Vmat1-T/T
Prrt3
Polb
Fst
Eif2a
Stx1a
RNG105
Ts1Cje
Actl6b
% Immobility in FS test
r = 0.25, P = 0.20
Vmat1-I/I
Begain
Vmat1-T/I
Il1r1
B2m
Aldh3a2
Caskin1
nSP
Scop
TMEM115
Prrt3
GluA1
Cntn4
RNG105
Fst
Ts1Cje
Actl6b
GT1
Papst1
LINC643
Trpm2
Vmat1-T/T
Eif2d
Act
Polb
Stx1a
Eif2a

**Figure S4. Correlations of brain lactate levels and pH with behavioral measures in a confirmatory cohort.** Scatter plots showing effect size-based correlations between actual lactate levels and pH, and behavioral measures. Data from 27 strains/conditions of mice used in the prediction analysis are shown. EP, elevated-plus maze; FS, forced swim test; LD, light/dark transition test; OF, open field test; *r*, Pearson's correlation coefficient.

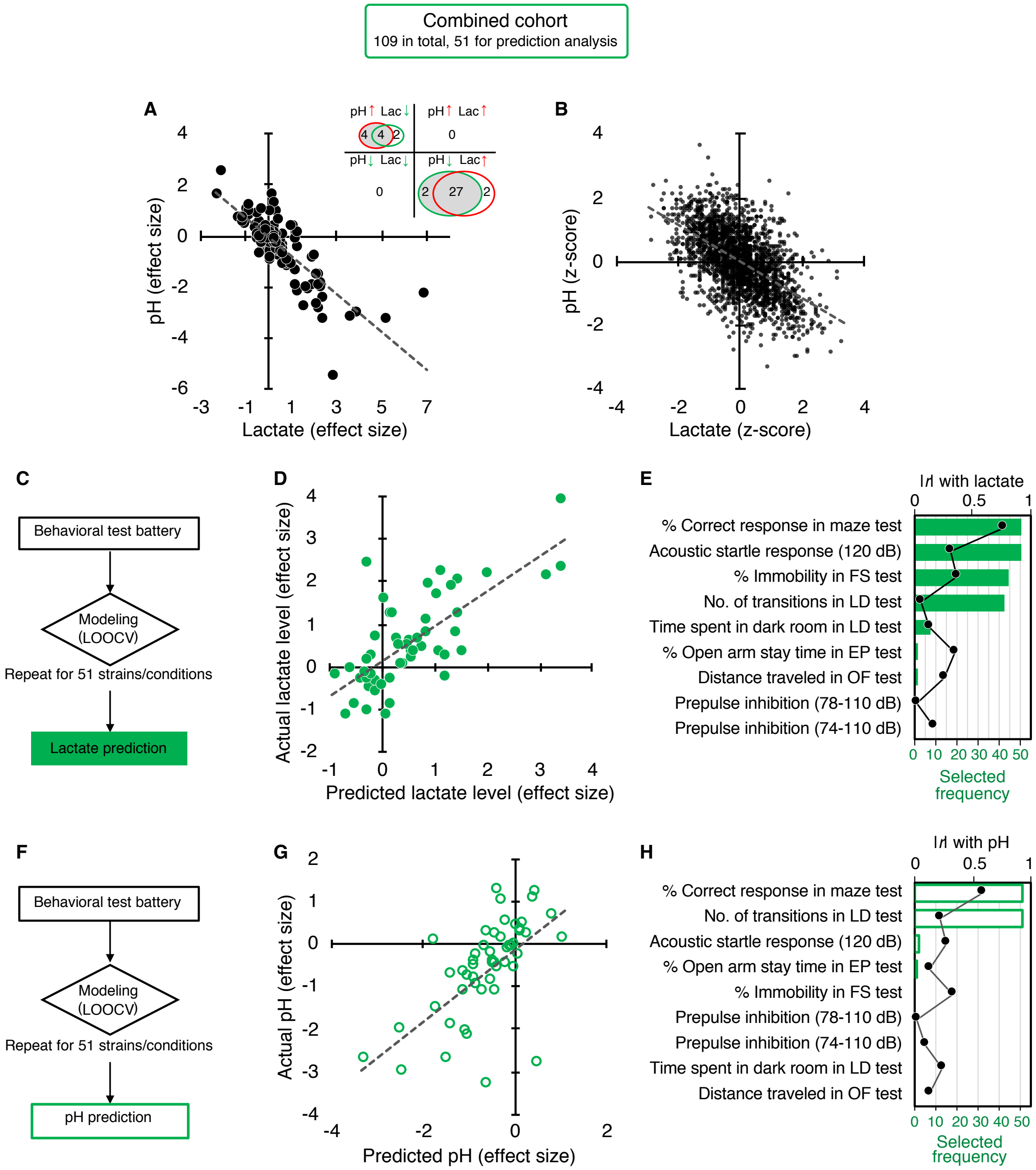


**Figure S5. Correlation of increased brain lactate levels and decreased pH and their associations with poor working memory: studies in a combined cohort.** (A) Venn diagrams show the number of strains/conditions of animal models with significant changes (P < 0.05 compared to the corresponding controls) in brain pH and lactate levels in a combined cohort. Scatter plot shows the effect size-based correlations between pH and lactate levels of 109 strains/conditions of animals combined. (B) Scatter plot showing z-score-based correlations between pH and lactate levels of 2,294 animals combined. A z-score was calculated for each animal within strain/condition. (C–H) Prediction of brain lactate levels (C–E) and pH (F–H) from behavioral outcomes in 51 strains/conditions of animals. The scatter plot shows correlations between predicted and actual lactate levels (D) and pH values (G). Feature preference for constructing the model to predict brain lactate levels (E) and pH (H). Bar graphs indicate the selected frequency of behavioral indices in the LOOCV. Line graph shows the absolute correlation coefficient between brain lactate levels and pH, and each behavioral index of 51 mouse strains. *r*, Pearson's correlation coefficient.

Lactate vs. behaviors

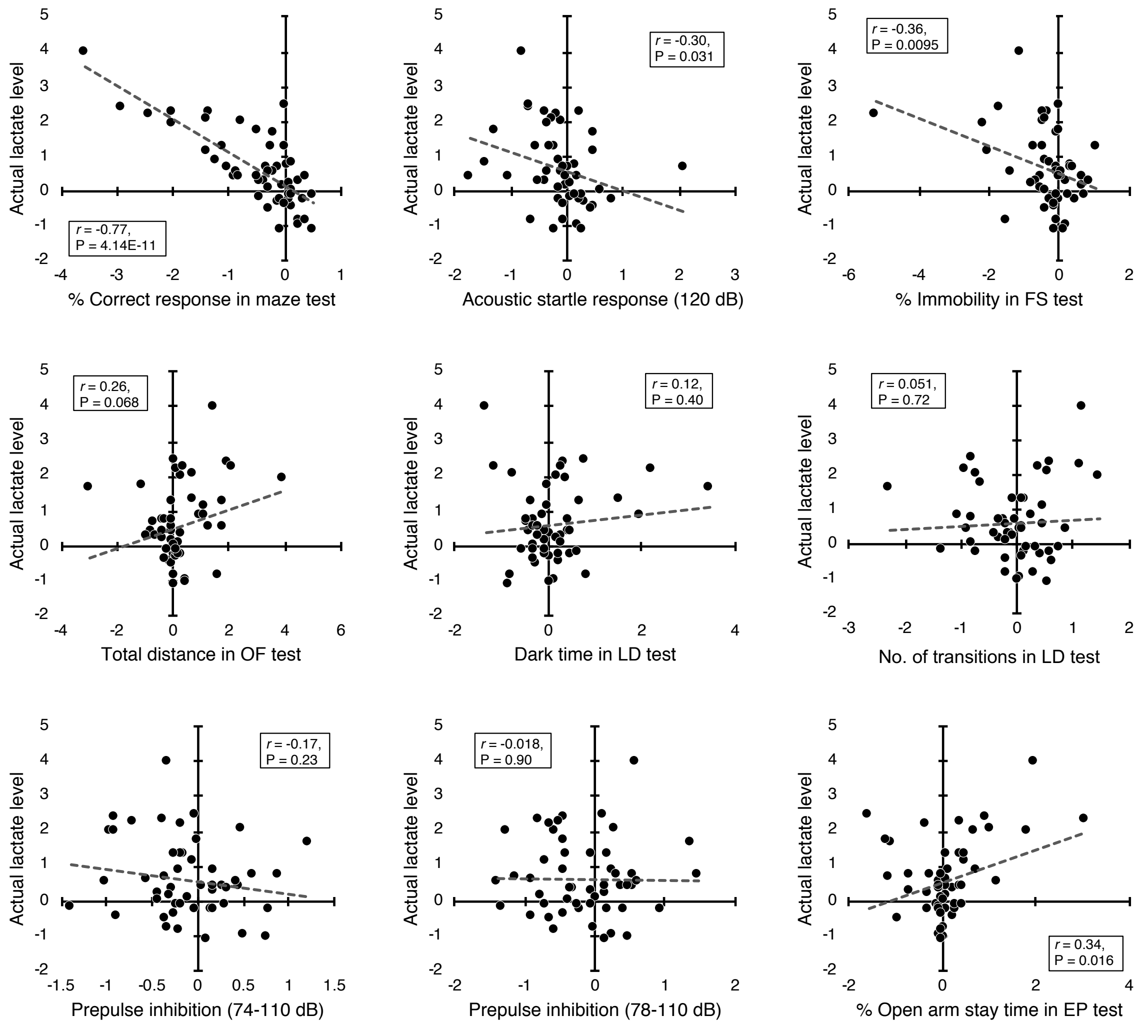


Continued

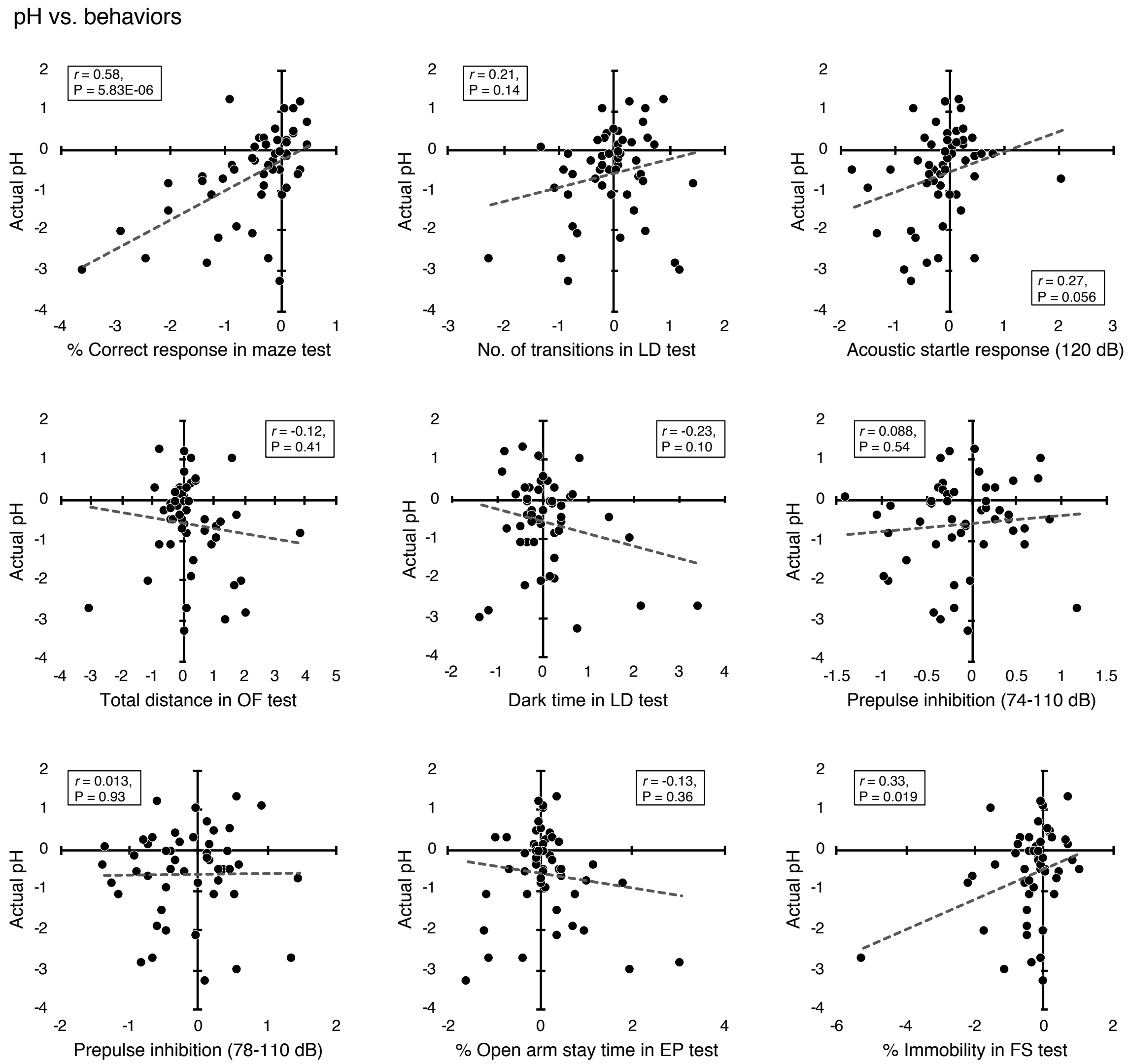


**Figure S6. Correlations of brain lactate levels and pH with behavioral measures in a combined cohort.** Scatter plots showing effect size-based correlations between actual lactate levels and pH, and behavioral measures. Data from 51 strains/conditions of mice used in the prediction analysis are shown. EP, elevated-plus maze; FS, forced swim test; LD, light/dark transition test; OF, open field test; *r*, Pearson's correlation coefficient.

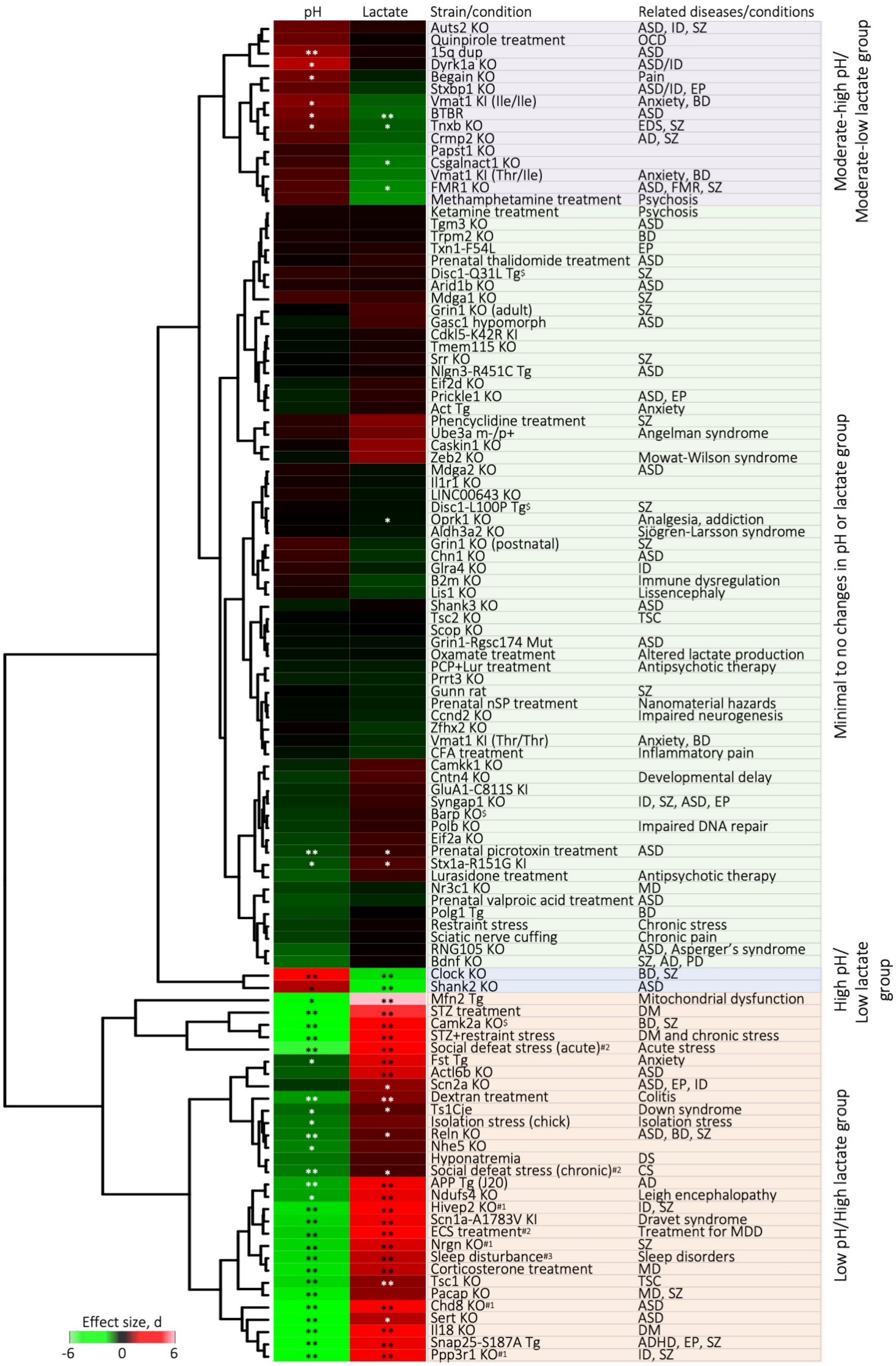
pH
Lactate
Strain/condition
Related diseases/conditions
Moderate-high pH/ Moderate-low lactate group
Minimal to no changes in pH or lactate group
High pH/ Low lactate group
Low pH/High lactate group
Effect size, d
-6
0
6
Auts2 KO
ASD, ID, SZ
Quinpirole treatment
OCD
15q dup
ASD
Dyrk1a KO
ASD/ID
Begain KO
Pain
Stxbp1 KO
ASD/ID, EP
Vmat1 KI (Ile/Ile)
Anxiety, BD
BTBR
ASD
Tnxb KO
EDS, SZ
Crmp2 KO
AD, SZ
Papst1 KO
Csgalnact1 KO
Vmat1 KI (Thr/Ile)
Anxiety, BD
FMR1 KO
ASD, FMR, SZ
Methamphetamine treatment
Psychosis
Ketamine treatment
Psychosis
Tgm3 KO
ASD
Trpm2 KO
BD
Txn1-F54L
EP
Prenatal thalidomide treatment
ASD
Disc1-Q31L Tg$
SZ
Arid1b KO
ASD
Mdga1 KO
SZ
Grin1 KO (adult)
SZ
Gasc1 hypomorph
ASD
Cdkl5-K42R KI
Tmem115 KO
Srr KO
SZ
Nlgn3-R451C Tg
ASD
Eif2d KO
Prickle1 KO
ASD, EP
Act Tg
Anxiety
Phencyclidine treatment
SZ
Ube3a m-/p+
Angelman syndrome
Caskin1 KO
Zeb2 KO
Mowat-Wilson syndrome
Mdga2 KO
ASD
Il1r1 KO
LINC00643 KO
Disc1-L100P Tg$
SZ
Oprk1 KO
Analgesia, addiction
Aldh3a2 KO
Sjögren-Larsson syndrome
Grin1 KO (postnatal)
SZ
Chn1 KO
ASD
Glra4 KO
ID
B2m KO
Immune dysregulation
Lis1 KO
Lissencephaly
Shank3 KO
ASD
Tsc2 KO
TSC
Scop KO
Grin1-Rgsc174 Mut
ASD
Oxamate treatment
Altered lactate production
PCP+Lur treatment
Antipsychotic therapy
Prrt3 KO
Gunn rat
SZ
Prenatal nSP treatment
Nanomaterial hazards
Ccnd2 KO
Impaired neurogenesis
Zfhx2 KO
Vmat1 KI (Thr/Thr)
Anxiety, BD
CFA treatment
Inflammatory pain
Camkk1 KO
Cntn4 KO
Developmental delay
GluA1-C811S KI
Syngap1 KO
ID, SZ, ASD, EP
Barp KO$
Polb KO
Impaired DNA repair
Eif2a KO
Prenatal picrotoxin treatment
ASD
Stx1a-R151G KI
Lurasidone treatment
Antipsychotic therapy
Nr3c1 KO
MD
Prenatal valproic acid treatment
ASD
Polg1 Tg
BD
Restraint stress
Chronic stress
Sciatic nerve cuffing
Chronic pain
RNG105 KO
ASD, Asperger's syndrome
Bdnf KO
SZ, AD, PD
Clock KO
BD, SZ
Shank2 KO
ASD
Mfn2 Tg
Mitochondrial dysfunction
STZ treatment
DM
Camk2a KO$
BD, SZ
STZ+restraint stress
DM and chronic stress
Social defeat stress (acute)#2
Acute stress
Fst Tg
Anxiety
Actl6b KO
ASD
Scn2a KO
ASD, EP, ID
Dextran treatment
Colitis
Ts1Cje
Down syndrome
Isolation stress (chick)
Isolation stress
Reln KO
ASD, BD, SZ
Nhe5 KO
Hyponatremia
DS
Social defeat stress (chronic)#2
CS
APP Tg (J20)
AD
Ndufs4 KO
Leigh encephalopathy
Hivep2 KO#1
ID, SZ
Scn1a-A1783V KI
Dravet syndrome
ECS treatment#2
Treatment for MDD
Nrgn KO#1
SZ
Sleep disturbance#3
Sleep disorders
Corticosterone treatment
MD
Tsc1 KO
TSC
Pacap KO
MD, SZ
Chd8 KO#1
ASD
Sert KO
ASD
Il18 KO
DM
Snap25-S187A Tg
ADHD, EP, SZ
Ppp3r1 KO#1
ID, SZ

**Figure S7. Hierarchical clustering of 109 strains/conditions of animals with respect to brain pH and lactate levels.** The effect size was calculated for each strain/condition and was used in this analysis. [#1]pH and lactate data have been previously reported(1). [#2]Lactate data have been reported previously [70]. [#3]Lactate data have been submitted elsewhere. Asterisks indicate significant effects of genotype/condition. $^{*}P < 0.05$, $^{**}P < 0.01$; unpaired *t*-test, or one-way or two-way ANOVA followed by *post hoc* Tukey's multiple comparison test. Detailed statistical analysis is shown in Table S3. AD, Alzheimer's disease; ADHD, attention-deficit/hyperactivity disorder; ASD, autism spectrum disorders; BD, bipolar disorder; CS, chronic stress; DM, diabetes mellitus; EDS, Ehlers-Danlos syndrome; DS, depression symptom; EP, epilepsy; FMR, Fragile X mental retardation; ID, intellectual disability, KI, knock-in; KO, knock out; MD, major depressive disorder; OCD, obsessive-compulsive disorder; PD, Parkinson's disease; SZ, schizophrenia; Tg, transgenic; TSC, tuberous sclerosis complex.

**A**

| | Dependent variable: | | | |
|---|---|---|---|---|
| | pH | | Lactate | |
| Variable | ltl | P-value | ltl | P-value |
| Intercept | 1630 | < 0.0001 | 138.9 | < 0.0001 |
| Age | 0.2815 | 0.7784 | 0.6189 | 0.5361 |
| Sex | 0.4591 | 0.6463 | 0.8773 | 0.3806 |
| Storage duration | 2.464 | 0.0139 | 0.2065 | 0.8364 |

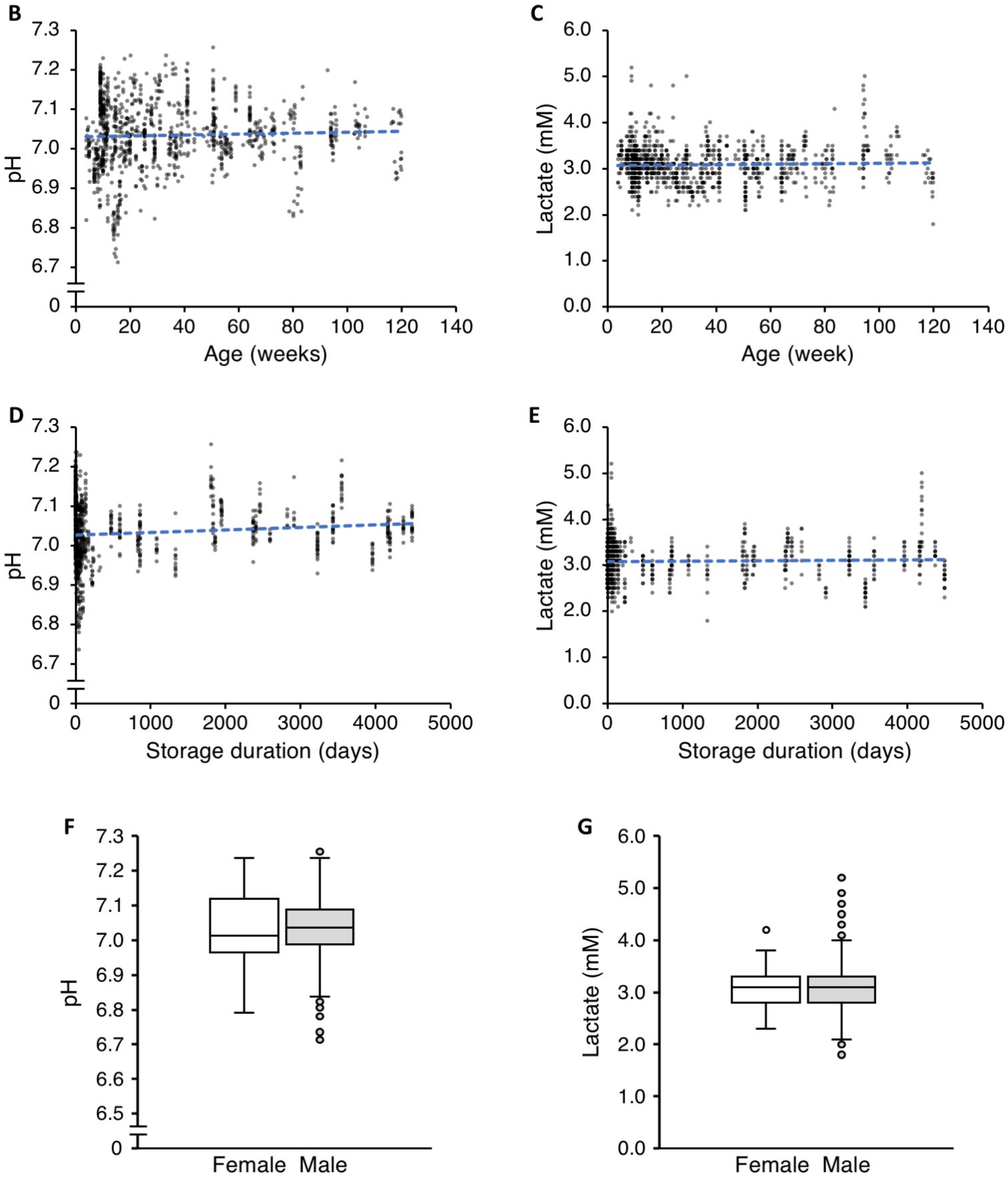

**Figure S8. Effects of age, sex, and storage duration on brain pH and lactate levels.** (A) Multivariate linear regression analysis. (B, C) Scatter plots showing correlations between age at sampling and raw pH (B), and lactate values (C) in wild-type/control animals. (D, E) Scatter plots showing correlations between storage duration and pH (D), and lactate values (E) in the wild-type/control animals. (F, G) Box plots of pH (F) and lactate values (G) in wild-type/control animals of each sex.
