## Supplementary Table 1 for "Large-scale animal model study uncovers altered brain pH and lactate levels as a transdiagnostic endophenotype of neuropsychiatric disorders involving cognitive impairment"

**Table S1.** Animal models used in this study

| | Name | Description | Related diseases, behaviors, or conditions |
|---|---|---|---|
| Exploratory cohort | | | |
| 1 | APP Tg | Mice expressing familial Alzheimer's disease-mutant human amyloid beta precursor protein (PDGF-hAPP$_{swe/Ind}$, line J20) (1) | AD (2,3) |
| 2 | Arid1b KO | Mice with heterozygous knockout of the AT-rich interaction domain 1b (4) | ASD (5,6) |
| 3 | Auts2 KO | Mice with heterozygous knockout of the Autism susceptibility candidate 2 (7) | ASD (8–10), ID (11), SZ (12) |
| 4 | Barp KO | Voltage gated calcium channel beta-anchoring and -regulatory protein KO mice (13) | Dysregulated voltage-gated $Ca^{2+}$ channel activity (13) |
| 5 | Bdnf KO | Mice with heterozygous knockout of the brain derived neurotrophic factor* (JAX stock #004339) | SZ, AD, PD (14) |
| 6 | BTBR | Inbred mouse strain BTBR T+ tf/J (15,16) | ASD (15,16) |
| 7 | Camk2a KO | Mice with heterozygous knockout of the calcium/calmodulin-dependent protein kinase II alpha (17–19) (JAX stock #002362) | BD (20–22), SZ (23) |
| 8 | Camkk1 KO | Mice with forebrain-specific constitutively active form of calcium/calmodulin kinase kinase 1 (24) | Impaired long-term memory (24) |

| | | | |
|---|---|---|---|
| 9 | Ccnd2 KO | Cyclin D2 KO mice (25) | Impaired adult brain neurogenesis (26,27) |
| 10 | CFA treatment | Mouse model of chronic inflammatory pain induced by complete Freund's adjuvant (CFA) (28,29) | Inflammatory pain (28,29) |
| 11 | Chd8 KO | Mice with heterozygous knockout of the long isoform of chromodomain helicase DNA-binding protein 8 (30) | ASD (31–35) |
| 12 | Chn1 KO | Chimerin 1 (α-chimerin) KO mice (36) | ASD (36) |
| 13 | Clock mutant | Mice with N-ethyl-N-nitrosourea-induced mutation in circadian locomotor output cycles kaput(37,38) (JAX stock #002923) | BD (39,40), SZ (41) |
| 14 | Corticosterone treatment | Mice chronically treated with corticosterone (42,43) | MD (44–46) |
| 15 | Crmp2 KO | Collapsin response mediator protein 2 KO mice (47) | AD (48), SZ (49) |
| 16 | Dextran treatment | Mice treated with dextran sulfate sodium (50) | Colitis (50) |
| 17 | Disc1-L100P mutant | Mice with N-ethyl-N-nitrosourea-induced L100P amino acid exchange mutation in exon 2 of Disrupted-in-Schizophrenia 1 (51) | SZ (52–54) |
| 18 | Disc1-Q31L mutant | Mice with N-ethyl-N-nitrosourea-induced Q31L amino acid exchange mutation in exon 2 of Disrupted-in-Schizophrenia 1 (51) | SZ (52–54) |
| 19 | Dyrk1a KO | Mice with heterozygous knockout of the dual specificity tyrosine phosphorylation regulated kinase 1a (55) | ASD/ID (6,56,57) |
| 20 | ECS treatment | Mice treated with electroconvulsive stimulation (58,59) | Treatment for MD (60,61) |
| 21 | Fmr1 KO | Fragile X mental retardation protein translational regulator 1 KO mice (62) | ASD, FMR, SZ (23) |
| 22 | Gasc1 hypomorph | Gene amplified in squamous cell carcinoma 1 hypomorphic mutant mice (63,64) | ASD (65) |
| 23 | Glra4 KO | Glycine receptor alpha 4 KO mice (66,67) | ID (68) |
| 24 | Grin1 KO (postnatal) | GABArgic neuron-specific glutamate receptor, ionotropic, NMDA1 KO mice | SZ (70,71) |

| | | | |
|---|---|---|---|
| | | (Protein phosphatase 1, regulatory subunit 2-cre; Grin1$^{loxP/loxP}$) (69) | |
| 25 | Grin1 KO (adult) | GABArgic neuron-specific glutamate receptor, ionotropic, NMDA1 KO mice (Protein phosphatase 1, regulatory subunit 2-cre; Grin1$^{loxP/loxP}$) (69) | SZ (70,71) |
| 26 | Gunn rat | Gunn rats (Gunn/Slc-j/j) (72) | SZ (73) |
| 27 | Hivep2 KO | Human immunodeficiency virus type 1 enhancer binding protein 2 (Schnurri-2) KO mice (74) | ID (75,76), SZ (74) |
| 28 | Hyponatremia | Mice treated with 1-deamino-8-D-arginine vasopressin and fed with a liquid formula (77–79) | DS (80,81) |
| 29 | Il18 KO | Interleukin 18 KO mice (82,83) | DM (84,85) |
| 30 | Ketamine treatment | Mice treated with ketamine (86) | Psychosis (87) |
| 31 | Lurasidone treatment | Mice treated with lurasidone (88) | Atypical antipsychotic therapy (89,90) |
| 32 | Mdga1 KO | MAM domain containing glycosylphospatidylinositol anchor 1 KO mice (91) | SZ (92–94) |
| 33 | Mdga2 KO | Mice with heterozygous knockout of the MAM domain containing glycosylphospatidylinositol anchor 2 (95) | ASD (96,97) |
| 34 | Methamphetamine treatment | Mice treated with methamphetamine (98) | Psychosis (99) |
| 35 | Nhe5 KO | $Na^+/H^+$ exchanger 5 KO mice (100) | |
| 36 | Nlgn3-R451C KI | Mice with R451C amino acid exchange mutation in neuroligin 3 (15,101) | ASD (102,103) |
| 37 | Nr3c1 Tg | Mice overexpressing glucocorticoid receptor under the Camk2a promoter[#] | MD (104) |
| 38 | Nrgn KO | Neurogranin KO mice (105–107) | SZ (108,109) |
| 39 | Oxamate treatment | Mice treated with sodium oxamate, an inhibitor of lactate dehydrogenase | Inhibition of lactate dehydrogenase |

| | | | |
|---|---|---|---|
| 40 | Pacap KO | Pituitary adenylate cyclase-activating polypeptide KO mice (110) | MD (111), SZ (112) |
| 41 | 15q dup | Mice with a paternal duplication of human chromosome 15q11-13 (113) | ASD (114–117) |
| 42 | Phencyclidine treatment | Subchronic phencyclidine-treated mice (88,118) | SZ (119) |
| 43 | PCP+Lur | Phencyclidine (PCP)- and lurasidone (Lur)-treated mice (88,118) | Atypical antipsychotic therapy (89,90) |
| 44 | Polg1 Tg | Forebrain-specific catalytic subunit of mitochondrial DNA polymerase KO mice (120) | BD (121) |
| 45 | Ppp3r1 KO | Forebrain-specific protein phosphatase 3, regulatory subunit B, alpha isoform (calcineurin B, type 1) KO mice (122–124) | SZ (125) |
| 46 | Quinpirole treatment | Mice treated with quinpirole, a dopamine D2 receptor agonist (126) | OCD (127) |
| 47 | Reln Tg | Mice lacking the C-terminal region of Reelin (128) | ASD (129–131), BD (132), SZ (133) |
| 48 | Restraint stress | Mice exposed to chronic restraint stress (134) | CS |
| 49 | Sciatic nerve cuffing | The sciatic nerve cuffing mouse model of neuropathic pain (135,136) | Chronic pain |
| 50 | Scn2a KO | Mice with heterozygous knockout of the sodium voltage-gated channel alpha subunit 2 (137) | ASD (138,139), EP (140–142), ID (143,144) |
| 51 | Sert KO | Serotonin transporter KO mice (145) | ASD (146,147) |
| 52 | Shank2 KO | SH3 and multiple ankyrin repeat domain 2 KO mice (148) | ASD (97) |
| 53 | Shank3 KO | SH3 and multiple ankyrin repeat domain 3b KO mice (149) (JAX stock #017688) | ASD (150–152) |
| 54 | Snap25-S187A KI | Mice with S187A amino acid exchange mutation in synaptosomal-associated protein of 25 kDa (153,154) | ADHD (155–160), EP (161,162), SZ (163,164) |
| 55 | Social defeat stress (acute) | Mice exposed to social defeat stress (165,166) | Acute stress |

| | | | |
|---|---|---|---|
| 56 | Social defeat stress (chronic) | Mice exposed to social defeat stress (167,168) | CS |
| 57 | STZ treatment | Mice treated with streptozotocin (169) | DM (170) |
| 58 | STZ + restraint stress | Mice treated with streptozotocin and exposed to chronic restraint stress (134,169) | DM and CS comorbidity (171) |
| 59 | Stxbp1 KO | Mice with heterozygous knockout of the syntaxin-binding protein 1 (172) | ASD/ID (57,144,173), EP (174,175) |
| 60 | Syngap1 KO | Mice with heterozygous knockout of the synaptic Ras GTPase-activating protein 1 (176,177) | ID, SZ, ASD (97), EP (174) |
| 61 | Thalidomide treatment | Rats prenatally exposed to thalidomide (178,179) | ASD (180) |
| 62 | Tnxb KO | Tenascin X KO mice (181,182) | EDS (183), SZ (184–186) |
| 63 | Tsc1 KO | Astrocyte-specific tuberous sclerosis complex 1 KO mice (Glial fibrillary acidic protein-cre; $Tsc1^{loxP/loxP}$) (187) | TSC (188) |
| 64 | Txn1-F54L Mut | Rats with a missense mutation in the thioredoxin 1 F51L generated by ENU-mutagenesis (189) | EP (189) |
| 65 | Valproic acid treatment | Mice prenatally exposed to valproic acid (190) | ASD (191) |
| Confirmatory cohort | | | |
| 66 | Act Tg | Mice with forebrain-specific overexpression of activin under the control of Camk2a promoter (192) | Anxiety-related behavior (192) |
| 67 | Actl6b KO | Actin-like 6B KO mice (193,194) | ASD (193) |
| 68 | Aldh3a2 KO | Aldehyde dehydrogenase 3 family member A2 KO mice (195,196) | Sjögren-Larsson syndrome (197) |
| 69 | B2m KO | Beta-2 microglobulin KO mice (JAX stock #002070) | Immune dysregulation (198) |
| 70 | Begain KO | Brain-enriched guanylate kinase-associated KO mice (199,200) | Neuropathic pain (199) |

| | | | |
|---|---|---|---|
| 71 | Caskin1 KO | Calcium/calmodulin-dependent serine protein kinase (CASK)-interacting protein 1 KO mice (201) | Enhanced nociception, anxiety-like behavior (201) |
| 72 | Cdkl5-K42R KI | Cyclin dependent kinase like 5 K42R knock-in mice# | |
| 73 | Cntn4 KO | Contactin-4 (BIG-2) KO mice (202) | Developmental delay (203) |
| 74 | Csgalnact1 KO | Chondroitin sulfate N-acetylgalactosaminyltransferase 1 KO mice (204,205) | Hyperactivity, increased acoustic startle response, altered social behavior (204) |
| 75 | Eif2a KO | Eukaryotic translation initiation factor 2A KO mice# | Deficiency of noncanonical translation initiation factors (206) |
| 76 | Eif2d KO | Mice with heterozygous knockout of the eukaryotic translation initiation factor 2D KO mice# | Deficiency of noncanonical translation initiation factors (206) |
| 77 | Fst Tg | Mice with forebrain-specific overexpression of follistatin under the control of Camk2a promoter (192) | Anxiety-related behavior (192) |
| 78 | GluA1-C811S KI | GluA1 palmitoylation-deficient (Cys811 to Ser substitution) knock-in mice (207,208) | Elevated seizure susceptibility, prolonged fear memory (207,208) |
| 79 | Grin1-Rgsc174 Mut | Mice with missense mutation in NMDA receptor subunit 1 (209) | Increased novelty-seeking behavior (209) |
| 80 | hMfn2-D210V Tg | Mice with pathogenic mutant of human Mitofusin 2 [hMFN2(D210V): Camk2a-tTA/TRE-hMFN2(D210V) Tg mice] (210,211) | Charcot–Marie–Tooth disease type 2A (212,213) |
| 81 | Il1r1 KO | Interleukin 1 receptor, type I KO mice (JAX stock #003245) | |

| | | | |
|---|---|---|---|
| 82 | Isolation stress | Layer-type neonatal chicks (*Gallus gallus*) exposed to isolation-induced stress (214,215) | Isolation stress |
| 83 | LINC00643 KO | Long intergenic non-protein coding RNA 643 (1700086L19Rik) KO mice[#] | |
| 84 | Lis1 KO | Mice with heterozygous knockout of the platelet-activating factor acetylhydrolase, isoform 1b, subunit 1 (216,217) | Lissencephaly (218,219) |
| 85 | Ndufs4 KO | NADH:ubiquinone oxidoreductase core subunit S4 KO mice (220) | Leigh encephalopathy (221,222) |
| 86 | nSP treatment | Prenatal treatment of amorphous nanosilica particle (223) | Reproductive and developmental hazards of nanomaterials |
| 87 | Oprk1 KO | Opioid receptor kappa 1 KO mice (224) | Analgesia, addiction |
| 88 | Papst1 KO | Mice with heterozygous knockout of the 3′-phosphoadenosine 5′-phosphosulfate (PAPS) transporter[#] | |
| 89 | Picrotoxin treatment | Mice prenatally exposed to $GABA_A$ receptor antagonist picrotoxin (225) | ASD (226) |
| 90 | Polb KO | Nex-Cre/DNA polymerase $beta^{fl/fl}$ mice (227) | Genetic diseases related to DNA repair (228,229) |
| 91 | Prickle1 KO | Mice with heterozygous knockout of the prickle1 (230,231) | EP (232,233), ASD (234,235) |
| 92 | Prrt3 KO | Actb-Cre/proline rich transmembrane protein $3^{fl/fl}$ mice (236) | Orphan metabotropic receptor |
| 93 | RNG105 KO | Mice with heterozygous knockout of the cell cycle associated protein 1 (237) | ASD, Asperger's syndrome (238) |
| 94 | Scn1a-A1783V KI | Mice carrying the sodium voltage-gated channel alpha subunit 1 A1783V mutation (239) | Dravet syndrome (240,241) |
| 95 | Scop KO | Suprachiasmatic nucleus circadian oscillatory protein KO mice (242) | Circadian regulation of memory formation (242) |

| | | | |
|---|---|---|---|
| 96 | Sleep disturbance | A mouse model of chronic sleep disorders caused by psychophysiological stress (243,244) | Sleep disorders (243,244) |
| 97 | Srr KO | Serine racemase KO mice (245,246) | SZ (247,248) |
| 98 | Stx1a-R151G KI | Syntaxin-1A (R151G) knock-in mice (249) | hyperactivity, anxiety- and depression-related behaviors, impaired working memory (249) |
| 99 | Tgm3 KO | Transglutaminase 3 KO mice (250) | ASD (251) |
| 100 | Tmem115 KO | Transmembrane protein 115 KO mice[#] | |
| 101 | Trpm2 KO | Transient receptor potential melastatin 2 KO mice (252) | BD (253,254) |
| 102 | Ts1Cje | Ts(16C-tel)1Cje mouse carrying a 7.6 Mb segmental trisomy of mouse chromosome 16 (255) | Down syndrome (256) |
| 103 | Tsc2 KO | Mice with heterozygous knockout of the tuberous sclerosis complex 2 (257–259) | TSC (188) |
| 104 | $Ube3a^{m-/p+}$ | Ubiquitin protein ligase E3A maternal-deficient mice (260) | Angelman syndrome (261) |
| 105 | Vmat1-Thr/Thr KI | Mice with humanized substitutions of the vesicular monoamine transporter 1 (replacement of the 133Asn of mouse Vmat1 with Thr or Ile) (262) | BD (263), Anxiety (264,265) |
| 106 | Vmat1-Thr/Ile KI | Mice with humanized substitutions of the vesicular monoamine transporter 1 (replacement of the 133Asn of mouse Vmat1 with Thr or Ile) (262) | BD (263), Anxiety (264,265) |
| 107 | Vmat1-Ile/Ile KI | Mice with humanized substitutions of the vesicular monoamine transporter 1 (replacement of the 133Asn of mouse Vmat1 with Thr or Ile) (262) | BD (263), Anxiety (264,265) |
| 108 | Zeb2 KO | *De novo* zinc finger E-box binding homeobox 2 Δex7/+ mice (266) | Mowat–Wilson syndrome (267,268) |

| | | | |
|---|---|---|---|
| 109 | Zfhx2 KO | Zinc finger homeobox 2 KO mice (269) | Hyperactivity, anxiety- and depression-related behaviors (269) |

AD, Alzheimer's disease; ADHD, attention-deficit/hyperactivity disorder; ASD, autism spectrum disorders; BD, bipolar disorder; CS, chronic stress; DM, diabetes mellitus; EDS, Ehlers-Danlos syndrome; DS, depression symptom; EP, epilepsy; FMR, Fragile X mental retardation; ID, intellectual disability, KI, knock-in; KO, knock out; MD, major depressive disorder; OCD, obsessive-compulsive disorder; PD, Parkinson's disease; SZ, schizophrenia; Tg, transgenic; TSC, tuberous sclerosis complex. *Mice with off-target deletion of conditional Bdnf allele derived from $Bdnf^{2lox}$ mouse line were used. #Unpublished mouse strain.
